# Evaluation of distal facial nerve branches contribution to facial nerve paralysis in rodents

**DOI:** 10.1101/2025.01.23.634588

**Authors:** Barbara G. Marin, Jordan L. Grant, Christine Mei, Giancarlo Tejeda, Ruby R. Taylor, Grant M. Lee, Victoria L. Ortega, Isabella G. Cozzone, Rachael M. Robbio, Trinity Robbio, Abhishek Prasad, Shekhar K. Gadkaree, Courtney M. Dumont

## Abstract

**Introduction/Aims:** Facial nerve paralysis is a complex and devastating condition. Translational research of facial paralysis recovery remains largely limited to animal studies, for which there are many potential models employed. When studying facial nerve regeneration in rodents, it is important to understand the converging contributions of the motor supply into the whisker pad. A consensus surgical approach and animal model has yet to be defined. Of particular interest for movement of the nose and whiskers are the buccal and marginal mandibular nerves. This study aims to evaluate how these distal nerve branches contribute to facial nerve paralysis and identify key morphological changes at the neuromuscular junctions (NMJs) in the whisker pad of rodents.

**Methods:** Adult rats underwent isolated transection of the buccal branch of the facial nerve, both the buccal and marginal mandibular branches of the facial nerve, or control sham surgery.

**Results:** Histological, electrophysiological, and behavior assessments confirmed that the transection of the buccal branch alone did not cease whisker movement in rats, but when combined with a transection of the marginal mandibular branch, it resulted in full paralysis of the whisker and nose movement.

**Discussion:** These results are indicative of the distinct roles of these nerves branches in facial paralysis repair following a transection injury. Further, our results suggest additional targets for facial nerve repair treatments.

## 1. Introduction

Facial paralysis is a devastating multifaceted condition for patients. The differential diagnosis for facial paralysis is broad-ranging, including idiopathic Bell’s palsy, Ramsay Hunt syndrome, Lyme disease, tumors, trauma, autoimmune disease, and more^1–4^. As a result of facial paralysis, an individual may suffer from the inability to blink, lose facial tone, and have brow or eyelid ptosis, nasolabial fold flattening, and hearing impairment. Thus, patients who have facial paralysis experience functional, physiological, social, and sociopsychological impairments^5–7^.

Management of facial paralysis remains complex, dependent on the mechanism of paralysis, timing of recovery, and severity of paralysis^8–11^. For primary transection, tension-free primary neurorrhaphy is the gold standard; however, it is often unachievable due to timing, size of lesion, and functional nerve availability^9^. If primary repair is not possible, autologous cable grafting remains the standard of care for facial nerve defects with gaps^9^. However, autografts rely on viable proximal and distal nerve endings. In the absence of a proximal nerve ending, facial reanimation using nerve transfers, static suspensions, or dynamic facial reanimation are all options. Despite advanced surgical methods, outcomes in patients with facial paralysis remain unsatisfactory. To date, surgical interventions are not always reliable in outcome, and moderate dysfunction post-treatment as well as continued experience of synkinesis can be a recurring problem^12,13^. Therefore, further development of treatment options for facial paralysis is crucial to significantly enhance patients’ quality of life^11,14–16^.

Treatment options for reconstruction of facial nerve injuries pose to be more challenging compared to other peripheral nerves due to the anatomical and etiological characteristics, such as size and convergence of innervation^17,18^. Rodent models of facial paralysis serve as a platform for testing new treatment options given their ability to achieve functional recovery and regrowth of motor nerves^19^. Rodent models of facial paralysis include nerve cuts, crush, and removal of segments to create gaps^20,21^. The rodent facial nerve preserves the anatomical architecture of the human facial nerve^20,22,23^. The site of injury for facial nerve models ranges from main trunk to the distal branches for implantation studies, specifically the buccal branch^24–31^. For treatments, including autografts or even more recent advances in biomaterials, the site of the main trunk cannot accommodate large gap injuries and treatment implantation in rodent models. This is in part due to the nerve retraction and anatomical variability between rodents. One key advantage of the distal nerve branch models is the location of the distal nerve branch, which prevents the retraction of the nerve into the skull base and the avoidance of dense vasculature that are encountered with main trunk injury models^32^.

For implantation studies, it is important to consider the convergence of innervation from both the buccal and marginal mandibular branch into the rodent whisker pad^33^. Despite the reports of the compensatory input of the marginal mandibular for whisker function, several studies only transect the buccal nerve for functional evaluation of recovery^34–42^. Given the model variability across research groups, the establishment of a reproducible distal facial nerve branch surgical technique is needed. Henstrom et al.^23^ provided a gross anatomical evaluation of the innervation to the follicles of the whisker pad. However, a detailed histological analysis of the neuromuscular junction (NMJ) to correlate their findings was not performed. In this study, we perform a direct comparison of electrophysiological, behavioral, and histological outcomes following two existing models of facial nerve injury. Evaluation of endogenous recovery over time in these disparate facial nerve injury models provides the foundation that may support a consensus being reached for studying mechanisms and therapeutic interventions following facial paralysis. The purpose of this study aims to evaluate how these distal nerve branches contribute to facial nerve paralysis and identify key morphological changes at the NMJ in the whisker pad of rodents.

## 2. Materials and Methods

### 2.1 Facial nerve injury model

Fourteen female Sprague Dawley rats (Envigo RMS, Indianapolis, IN) weighing 160 to 180 grams were used in all experiments. Animal experiments were performed with approval and in regulation with the Institutional Animal Care and Use Committee (IACUC) guidelines at the University of Miami. For all surgical interventions and electrophysiological studies, rats were anesthetized with 2% isoflurane in 1 L/min oxygen with depth of anesthesia confirmed *via* toe pinch and an ophthalmic lubricant was applied. A horizontal 10 mm incision was made from skin to the pre-parotid fascia. Following blunt dissection of the subcutaneous fascia, the parotid gland, buccal nerve, marginal mandibular nerve, parotid duct and the masseter muscle were exposed. The buccal and marginal mandibular nerves were traced to identify the convergence at the distal pes. The parotid duct was identified and separated from the marginal mandibular nerve to isolate the nerve. A 5-mm nerve gap defect was created in the left facial nerve branches using micro scissors to ensure complete transection and wound site was sutured. To evaluate the functional and behavioral outcomes across the facial nerve branches, rats were divided into three groups (n = 4-5 per group) (sham surgery, buccal transection (BB), buccal + marginal mandibular transection (BB+MM) (Fig.1). Pain and inflammation were managed with 0.5 mg/kg carprofen. The right facial nerves remained intact, serving as an uninjured control for each animal. Animals were monitored to assess recovery following anesthesia, after which they were assessed daily for 7 days to ensure resumption of normal eating, drinking, and grooming, and then monitored weekly.

**Figure 1.**
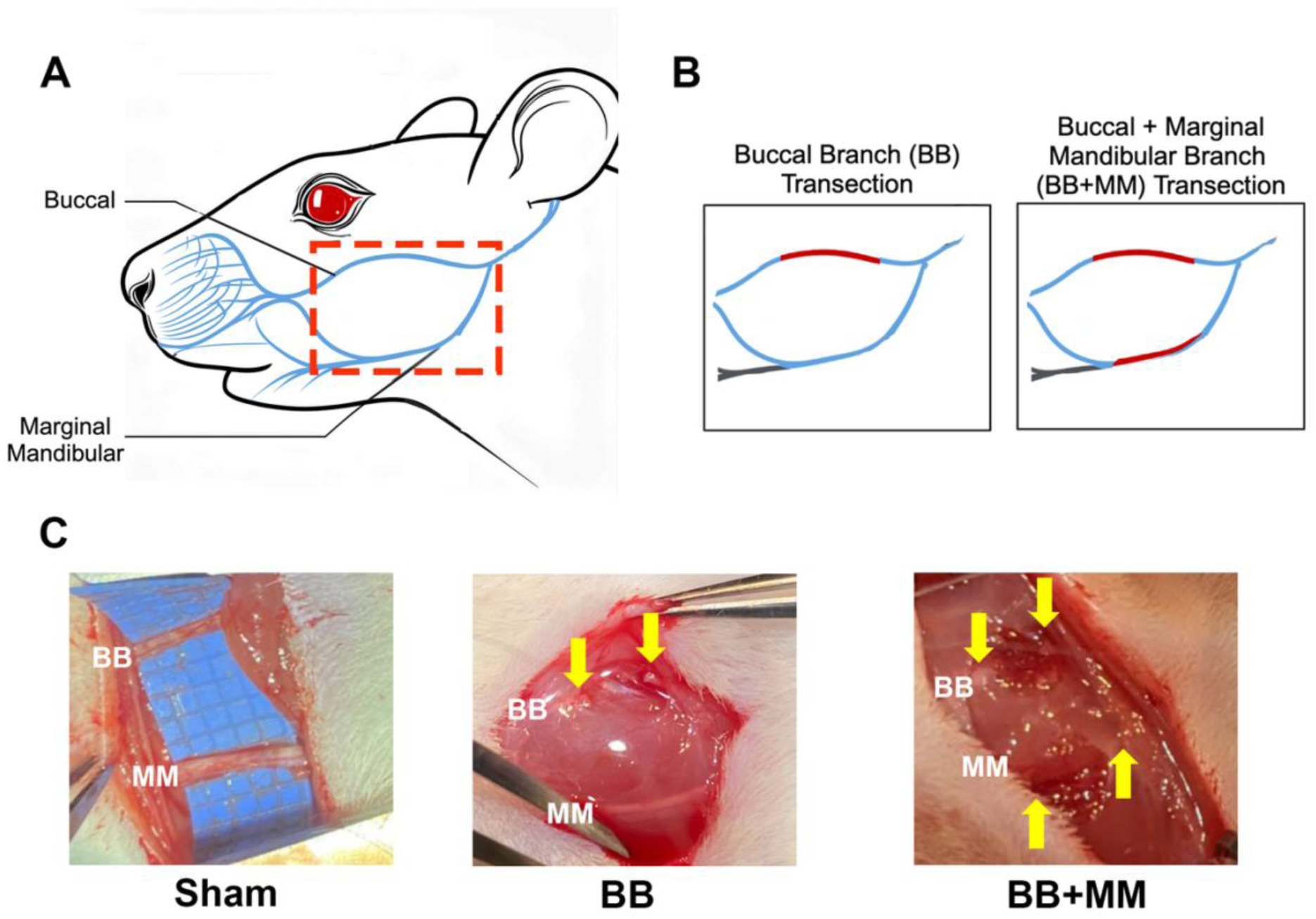
Overview of facial nerve surgical models. A) Diagram of rat facial nerves indicating main trunk branches that innervate the rodent nose and whisker pad muscles. B) Facial nerve transection models, buccal branch only 5 mm transection, and buccal branch + marginal mandibular branch 5 mm transection. C) Surgical photographs showing the following conditions: sham (both branches intact), buccal branch only transection (BB) and marginal mandibular intact, buccal branch + marginal mandibular branch (BB+MM) transection. BB, buccal branch; MM, marginal mandibular branch of the facial nerve.

### 2.2 Facial Palsy Assessment

Animal facial palsy was assessed on a scale of 0-7, with points accumulated for whisker and nose symmetry as established previously^43^. A seven-point indicated a normal midface, and a zero-point indicated complete facial palsy. Facial palsy score (FPS) was independently evaluated by three blinded individuals and were carried out weekly for 8 weeks post-injury. The distal branches such as the buccal branch allows for measurable behavioral outcomes due to its innervation into the whisker and nose musculature, which encompass the foundation of the FPS in rodents. For implantation studies, this approach is chosen partly because accessing the main trunk in small rodent models is technically challenging, which can reduce model reproducibility.

### 2.3 Compound Muscle Action Potential Recordings of the Mystacial Pad Muscles

Electrophysiological recordings were performed from the mystacial pad muscles 8 weeks after injury similar to previous reports^44^. A mid-sagittal incision was performed to expose the rat skull and to locate the bregma. Two stainless steel screws (3/16 inch, 0– 80, CDE Fasteners, NJ) were manually drilled into the skull on diagonal quadrants for separate grounding of the stimulating and recording electrodes. The buccal and marginal mandibular nerves were exposed distal to the condition for stereotactic 1-mm insertion of the stimulating electrode, which was a single 1-mm long and 250-µm diameter tungsten electrode (A-M Systems #563410, WA). The whisker pad was exposed and a 16-channel microwire electrode array (MEA) (Tucker-Davis Technologies (TDT), Alachua, FL), used as the recording electrode, was inserted 2-mm into the whisker pad with a hydraulic micro positioner (FHC, VT).

Stimulation was delivered independently to the buccal or marginal mandibular nerve with an isolated pulse stimulator (A-M Systems, Model 2100, Sequim, WA). A 100- µs duration, 20-ms pulse width, and 1-ms pre-train delay of single train, biphasic, cathode leading, symmetric square wave was applied. Stimulation currents of increasing increments of amplitude (1mA, 2mA, and 4mA) were delivered in a successive pulse train for 1-min each with a 3-min rest period in between each train. The 2mA current was determined as the suprathreshold intensity. Evoked compound muscle action potentials (CMAPs) were recorded by the MEA at the whisker pad at 6-kHz sampling rate with a pre-amplifier (PZ5, TDT) and a digital signal processor (RZ2, TDT) that converted the analog into digital signals.

Data analysis of the evoked electrophysiological responses was performed in Matlab (Mathworks, MA). A high-pass filter of 1-Hz cutoff frequency was used to remove the baseline drift and low frequency artifacts from the data. The medium CMAP response from the 2mA stimulation session was used for all analysis^44^. The amplitude, latency, and duration of the first 50 CMAPs for each animal were averaged and used for analysis, approximately corresponding to the first second of stimulation. The CMAP amplitude, latency, and duration of the response was calculated within a 5-ms window following the onset of stimulation artifact^44^. For injury conditions in which CMAPs were not evoked by any amount of stimulus current, the response was quantified as 0mV for amplitude 0-ms for the duration of the response. A maximum latency of 5-ms was assigned to match the analysis window which suggested that a response was not registered during this window.

### 2.4 Identification of penetrating axons

Buccal branches of the facial nerve were flash frozen, and cryosectioned (Leica, Deer Park, IL) longitudinally in 10µm sections at 8 week following injury. Sections were fixed, permeabilized as necessary, and incubated overnight at 4°C with the following primary antibodies: rabbit anti-neurofilament-200 (1:200, No. N4142, Sigma-Aldrich, St. Louis, MO), mouse anti-S100 calcium-binding protein B (S100β, 1:250, Sigma-Aldrich No. S2532-100µL). Species-specific fluorescent secondary antibodies were used for identification at 1:1000 (Thermo Fisher, Carlsbad, California). Following secondary, nerve slices were incubated with red-fluorescent myelin marker FluoroMyelin^TM^ (1:150, Thermo Fisher Scientific, Waltham, MA, No. F34652). Hoechst 33342 (1:2000, Life Technologies, Carlsbad, CA) was used as a counterstain for all tissue sections. Tissues were imaged on a fluorescent microscope (Eclipse Ti Series, Nikon Instruments, Melville NY) and were processed in ImageJ (NIH, Bethesda, MD, USA).

### 2.5 Characterization of muscle innervation and neuromuscular junction

After 8 weeks post injury, whisker pad muscle beds were flash frozen, and cryosectioned (Leica) at 20μm thickness. Muscle sections were fixed, permeabilized as necessary, and incubated overnight at 4°C with the following primary antibodies: rabbit anti-neurofilament-200 (1:200) and mouse anti-synaptic vesicle 2 (SV2, 1:200; Developmental Studies Hybridoma Bank Ab2315387, Iowa City, IA). Species-specific secondary antibodies were used at 1:1000 (Invitrogen no. AB150077). Following secondary, sections were incubated with rhodamine-conjugated α-bungarotoxin binding to the acetylcholine receptor (AChR) (1:50; Thermo Fisher Scientific no. B35450). All sections were counterstained with Hoechst 33342 (1:2000).

Images were obtained on a Dragonfly Spinning Disk Confocal (Andor Technologies, Belfast, Ireland) using 1024 x 1024 frame size, 63x objective, 1.5x zoom, and 1 μm z-stack interval. Only *en face* NMJs were analyzed. There were six individual muscle tissues that were analyzed per animal per condition (sham: 100 NMJs; BB: 98 NMJs; BB+MM: 57 NMJs). Presynaptic innervation was determined based on the presence of axon (NF200+) and presynaptic innervation (SV2+) in the post synaptic endplate (AChR+). The synapses were further analyzed in ImageJ using the NMJ-morph plug-in for evaluating pre and post synaptic components of the NMJs^45^.

### 2.6 Statistical analysis

All results were analyzed while blinded to treatment and surgical injury paradigm. The normality of residuals was analyzed for each data set using the D’Agostino-Pearson omnibus test as well as visual inspection of the QQ plots. For the FPS a two-way ANOVA with the Geisser-Greenhouse correction tests followed by a Tukey’s multiple comparison test was performed. CMAP assessments were analyzed by one-way ANOVA with Tukey post hoc for multiple comparisons. For NMJ, morphological metrics were analyzed by a one-way ANOVA followed by a Tukey post hoc multiple comparison. Prism 10 (GraphPad Software, La Jolla, CA) software was utilized for all data analysis. Statistical significance was determined at *p* < 0.05. All data are represented as mean values ± SEM unless otherwise stated.

## 3. Results

### 3.1 Buccal transection alone is insufficient for establishing prolonged facial palsy

Animals underwent nerve transections on the left side of their face and the right side served as the contralateral control for evaluation of functional recovery (Fig. 2A). Scores were evaluated on a scale from 0 indicating full facial paralysis and 7 indicating normal facial movement. Functional facial recovery was assessed weekly post-transection for 8 weeks (Fig. 2B). The buccal and marginal mandibular transection group (BB+MM) had significantly lower overall FPS than the buccal only (BB) transection group and sham group from injury throughout the 8 weeks of assessment. There was a significant difference between the sham and BB group between 1- and 3-weeks following injury. However, after 3 weeks the overall FPS was not significantly different between the sham and the BB transection group suggesting rapid recovery of function after transection of the buccal branch only (Fig. 2C).

**Figure 2.**
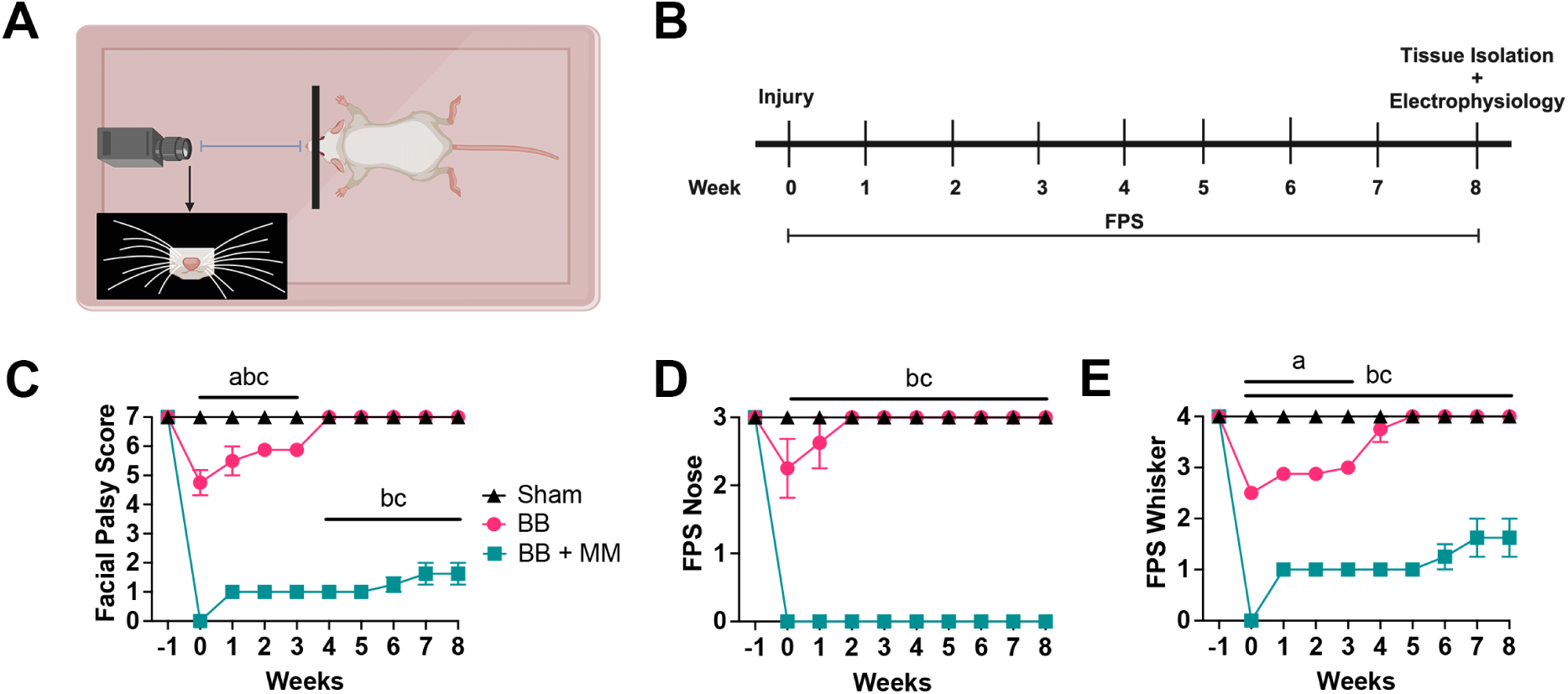
Functional assessment of facial palsy recovery. A) Schematic of rodent facial palsy score (FPS) functional assessment set up. B) Timeline of study from surgical transection of facial nerves at week 0, followed by weekly FPS assessment, and week 8 indicating tissue isolation and electrophysiological assessments. C) Overall FPS on a scale of 0-7, where points are accumulated for whisker and nose symmetry. FPS was assessed from the sham, the buccal branch only transection (BB), and the buccal branch and marginal mandibular transection group (BB+MM). Comparison between sham vs BB group a = p < 0.05; sham vs BB+MM, b = p < 0.0001; BB vs BB+MM, c = p < 0.0001. D) FPS subscores indicating nose symmetry sham vs BB+MM (b = p < 0.0001) and BB vs BB+MM (c = p < 0.0001). (E) Whisker symmetry FPS subscore sham vs BB (a = p < 00.05); sham vs BB+MM (b = p < 0.0001); BB vs BB+MM group (c = p < 0.05) from weeks 0-8. Data are represented as mean ± SEM, n=4 animals per condition. SEM, standard error of mean. (Figure create with BioRender).

Further evaluation of FPS via the individual subscore indicating nose symmetry resulted in a significantly decreased score in the BB+MM group compared to sham and BB throughout the 8 weeks (Fig. 2D). However, the FPS nose subscore for the BB group was not significantly different than the sham group starting at only one week post injury. Similarly, the whisker symmetry FPS subscore was significantly decreased in the BB+MM group compared to sham and BB group from weeks 0-8. The BB group was significantly lower than the sham from week 0 – 3. However, the FPS whisker subscore for the BB group was not significantly different from the sham group after 3 weeks following injury (Fig. 2E). The points accumulated in the whisker panel for the BB+MM condition highlight the spasticity of minor trembling (Fig. 2E). The subscores of the FPS highlight the important complementary innervation of the marginal mandibular branch into both the rat nose and whisker muscles. It is imperative to transect both nerves to ensure full facial paralysis, especially if using behavioral assessments that evaluate the return of facial function.

### 3.2 Transection of buccal and marginal mandibular branches is necessary to eliminate electrophysiological signal conduction to the vibrissal musculature

Facial paralysis was also evaluated by electrophysiological activity, where either the buccal branch or the marginal mandibular were stimulated for all groups and resulting CMAPs were recorded and quantified. As shown by a representative trace in Figure 3C, when the buccal branch was stimulated the BB animals that did have evoked CMAP activity displayed an observable reduced amplitude, longer latency, and shorter duration. When quantified, as shown in Figure 3E, there was a reduction in CMAP amplitudes between the sham and BB only groups, and this difference was significant between the sham and BB+MM group. There was also a significant difference between sham and BB CMAP latencies and duration, and this was exacerbated between sham and BB+MM.

**Figure 3.**
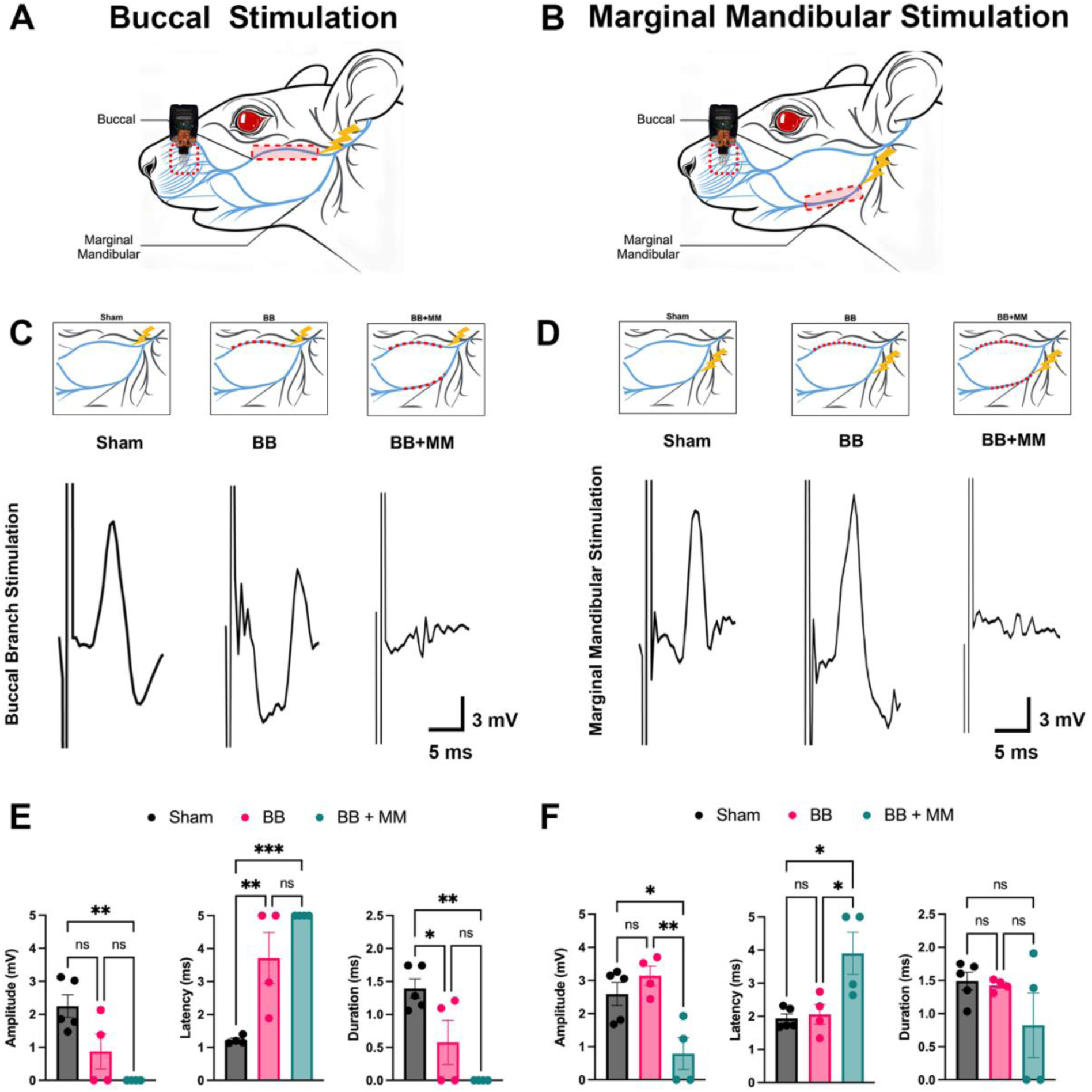
Electrophysiological analysis of compound muscle action potentials (CMAP) on buccal and marginal mandibular branches of sham, BB, and BB+MM transections at 8 weeks following injury. A) Diagram indicating stimulation of the buccal branch of facial nerve proximal to the injury site and TDT recording electrode on whisker pad of rodent. B) Diagram indicating stimulation of the marginal mandibular branch of facial nerve proximal to the injury site and TDT recording electrode on whisker pad of rodent. C) CMAP from whisker pad following stimulation of the buccal branch of the facial nerve in the sham, BB group, BB+MM group. D) CMAP from whisker pad following stimulation of the marginal mandibular branch of the facial nerve in the sham, BB group, and BB+MM group. E) Amplitude, latency and duration results of compounds muscle action potentials following buccal branch stimulation. p <0.05 *; p< 0.001 *** F) Amplitude, latency and duration results of compounds muscle action potentials following marginal mandibular branch stimulation. p < 0.05 *; p < 0.001 **. Data are represented as mean ± SEM, n= 4-5 animals per condition. SEM, standard error of mean.

As shown in Figure 3D, when the marginal mandibular was stimulated a CMAP was still observed in sham and BB animals as each of these groups had a healthy marginal mandibular nerve. When quantified, there was also no significant difference found between the sham and BB groups in CMAP amplitude, latency, or duration. This suggests that even if the BB is transected, there could still be electrophysiological activity in the whisker pad conducted by the marginal mandibular nerve. In the BB+MM group, CMAP amplitude was significantly decreased from sham and BB only and latency was significantly increased. There was also no significant difference in CMAP duration across all groups in marginal mandibular stimulation. These results suggest that transecting both the buccal and the marginal mandibular branch more effectively diminishes electrophysiological activity in the whisker pad.

### 3.3 Buccal nerve growth is limited at 8 weeks post-injury

Histological analysis of the regenerated nerves by immunofluorescence was subsequently performed at 8 weeks post-injury. A qualitative increase in cell infiltration was observed in BB and BB+MM conditions compared to sham controls (Fig 4B), which was expected due to rapid proliferation that occurs after nerve injury. Expression of the Schwann cell marker S100β was evident in all conditions, however, there was a marked decrease in S100β in the BB+MM condition. Further differences in nerve fascicle organization were observed with the expression of NF200 and fluoromyelin, markers for axons and myelin, respectively. While immunohistochemistry studies showed similar expression of Fluoromyelin and NF200 in the BB group and sham group, lower expression of these markers were observed in the BB+MM group.

**Figure 4.**
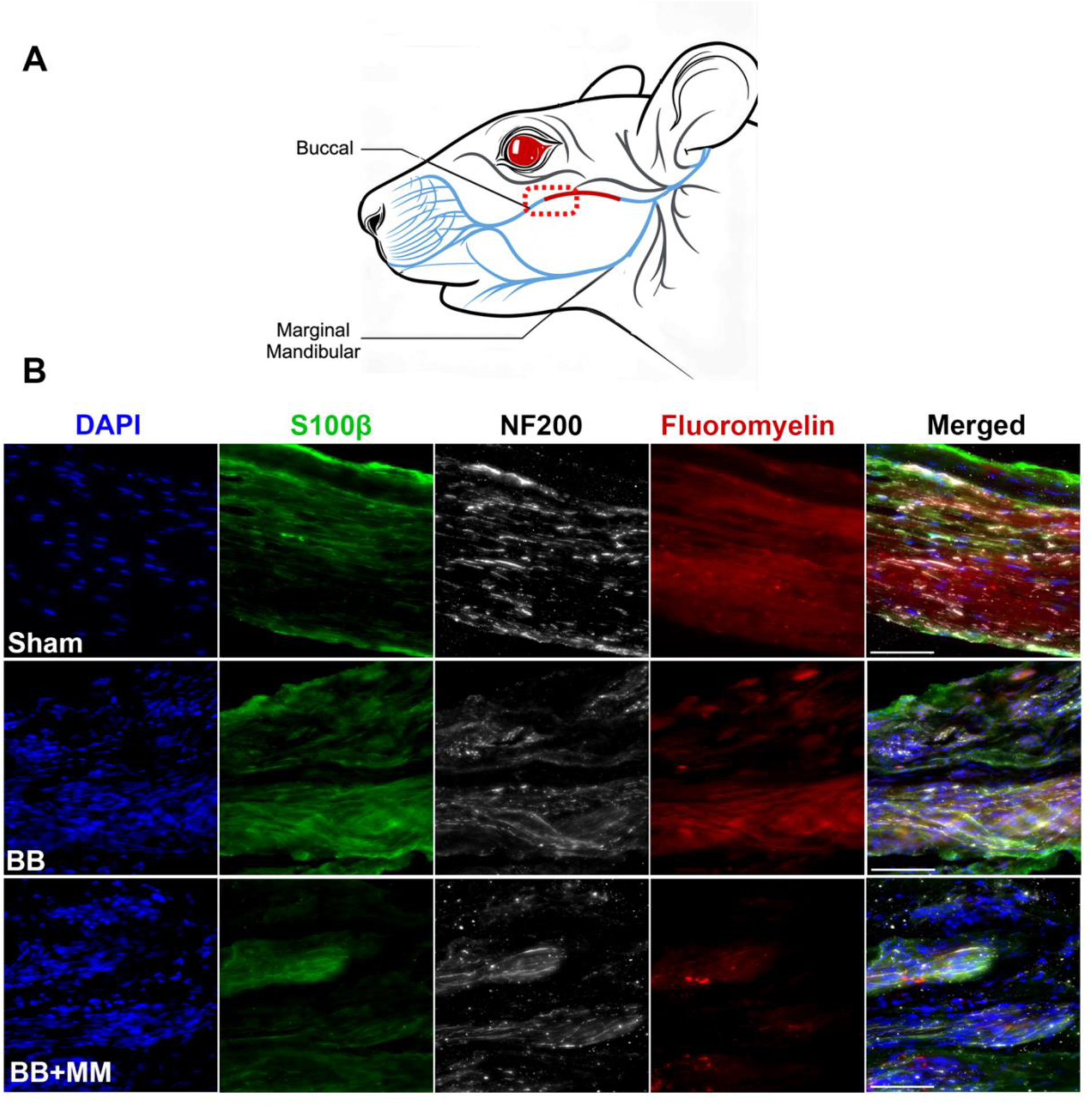
Histological analysis of regenerated rat facial nerve at 8 weeks post-injury. A) Diagram indicating the site of histological analysis in the buccal branch of the rodent facial nerve. B) Immunohistochemistry showed decreased expression of Fluoromyelin (red), S100β (green), and NF200 (grey) and disorganized alignment of nerves from the BB+MM as compared to the BB only and Sham group. Scale bar = 100μm. Cell nuclei were stained with DAPI (blue).

### 3.4 Transection of both buccal and marginal mandibular branches results in decreased neuromuscular innervation

Evaluation of NMJ morphology was performed at 8 weeks following injury. Muscle innervation was compared between the sham, BB, and BB + MM to evaluate the impact of the injury model. Innervation was determined by the absence or presence of both presynaptic NF200+ and SV2+ markers at the AchR+ motor endplate (Fig. 5A-B). There were no significant differences observed in the presynaptic NMJ morphology metrics of axon diameter and complexity across all conditions. There were significantly smaller branches in the BB+MM group compared to sham (Fig. 5C-D). Importantly, the total number of NMJs was significantly reduced in the BB + MM condition compared to the sham and BB condition (Fig. 5F). The BB group showed significantly higher numbers of innervated NMJs compared to the BB+MM group and not significantly different than the sham group. While the BB + MM group had significantly less innervated NMJs compared to the sham (Fig.5G). Overall, these results indicate that transection of both the buccal and marginal mandibular nerves results in fewer presynaptic elements, and of those remaining they are less likely to be innervated and/or have shorter branch length associated with NMJ disassembly.

**Figure 5:**
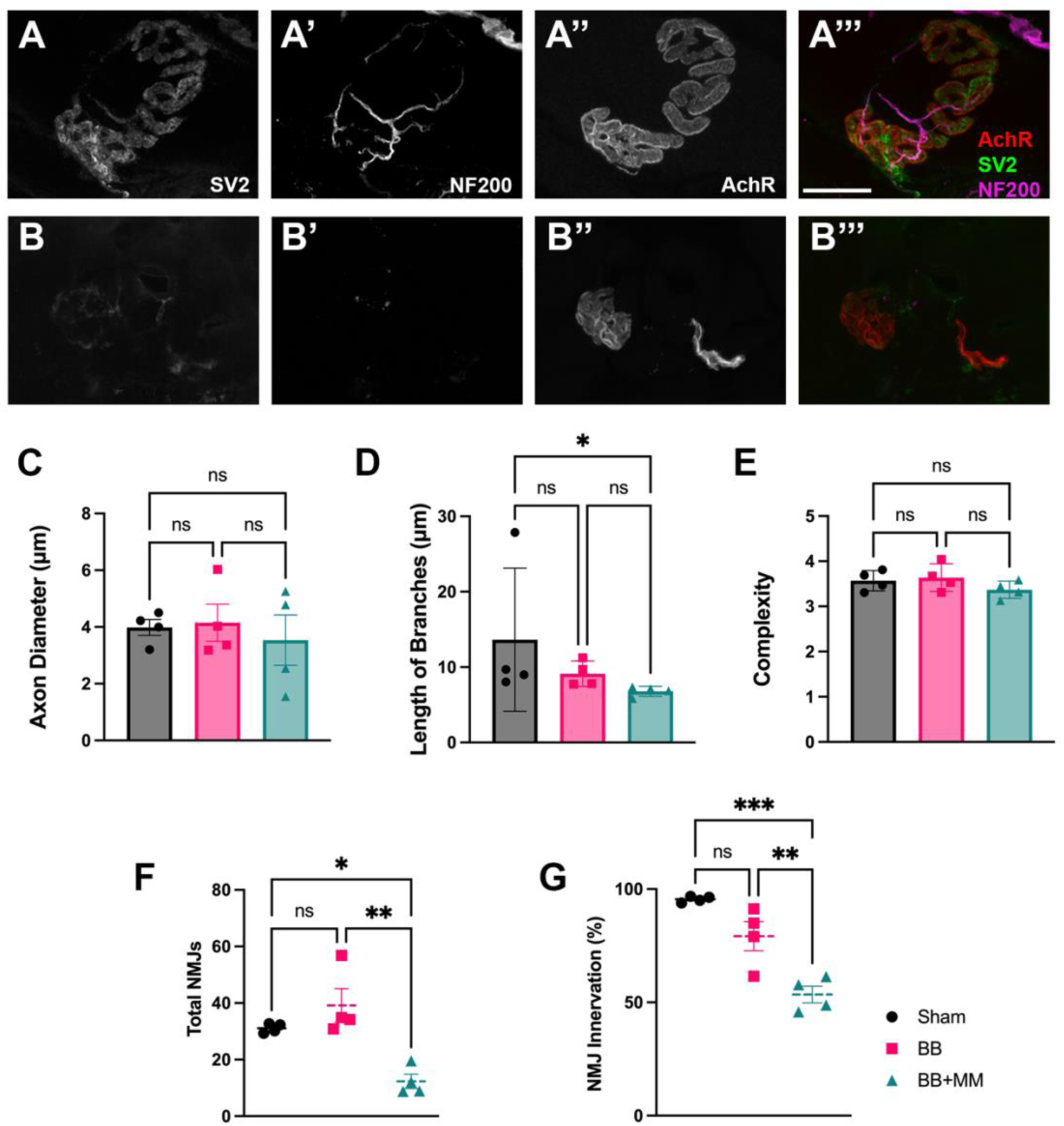
Neuromuscular innervation following facial nerve injury. Quantification of innervation of NMJs of sham, BB, and BB+MM transections at 8 weeks following injury. A-B Indicate images of NMJs. Axon and nerve terminal present within an innervated junction (A, A′, A′′, A′′′). Lack of a nerve terminal in a denervated junction (B, B′, B′′, B′′′). Scale bar = 25 μm (C-E) Presynaptic NMJ morphology variables for each group. (F) Total average of NMJs identified in the whisker pad of the rodents at 8 weeks post injury. NMJ innervation is indicated by presynaptic markers SV2 (green) and NF200 (magenta) within each motor endplate labeled with AChR (red). G) The standard error of mean of innervated NMJs are shown at 8 weeks post-injury. Data are represented as mean ± SEM, n= 3-4 animals per condition p < 0.05 *; p < 0.001 **; p < 0001***. SEM, standard error of mean.

Qualitative observations of the NMJs show marked differences per conditions. Concurrent with the innervation results, the BB + MM condition had significantly different morphology than the sham and BB groups in the motor endplate parameters generated by the NMJ-morph (Fig 6A). The NMJ morph pre-synaptic and post-synaptic morphology metrics demonstrate that the BB + MM group showed significantly greater differences compared to the other conditions (Fig 6B-I). The marked overall conclusion is that the BB group NMJs resembled the morphology to the NMJs in the sham group, whereas the BB+MM group were significantly reduced.

**Figure 6.**
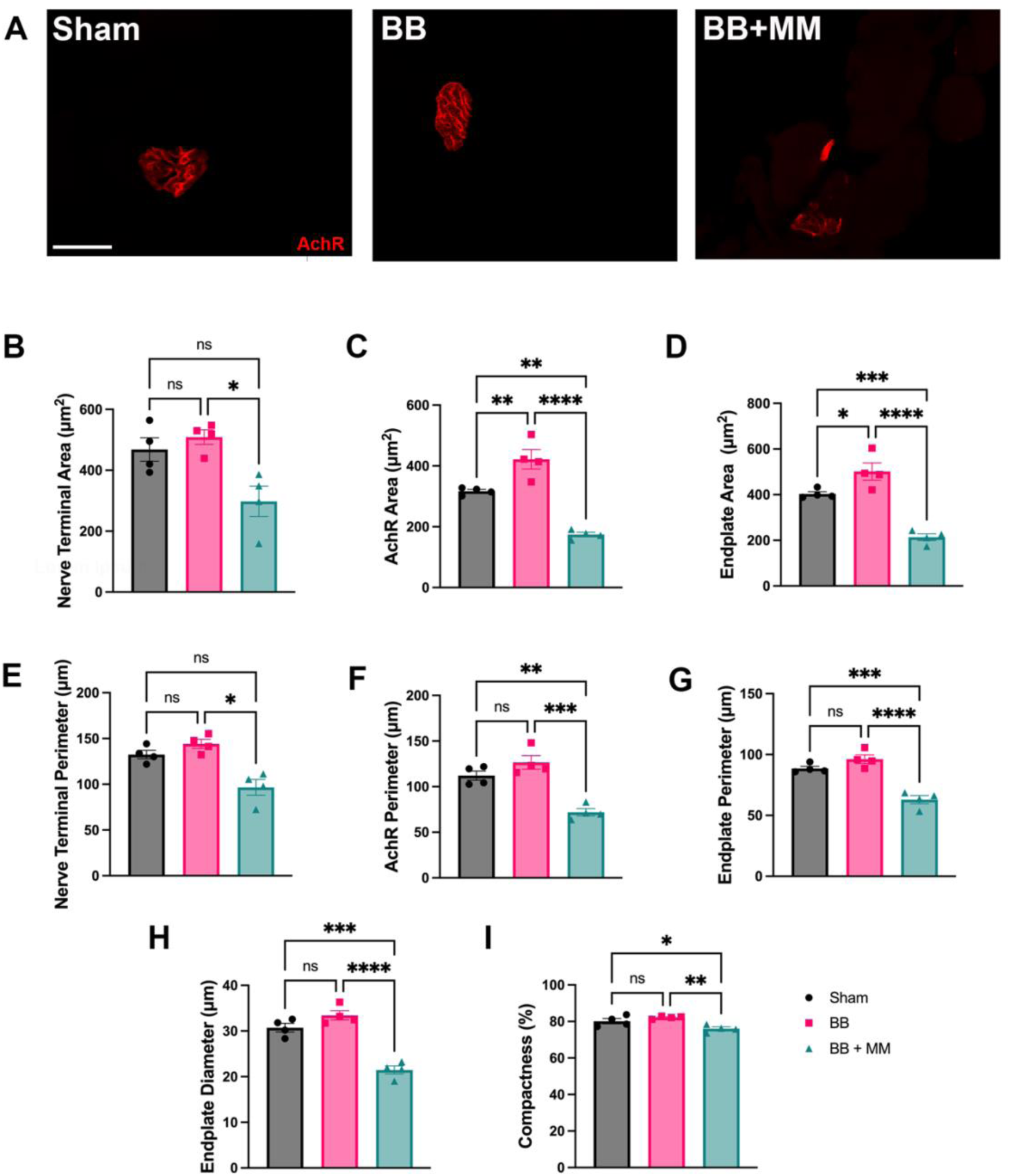
Effect of injury model on neuromuscular junction morphology at 8 weeks post-injury. A) Representative endplates from each group sham, BB only, BB + MM at 8 weeks post-injury. Scale bar = 25μm. B-I) Bar graphs demonstrating presynaptic and postsynaptic NMJ morphology variables. Data are represented as mean ± SEM, n= 4-5 animals per condition p < 0.05 *; p < 0.001 **; p < 0001***. SEM, standard error of mean.

## 4. Discussion

In this study, we evaluate endogenous recovery over time in two disparate facial nerve injury models to support the consensus and use of a double injury distal branch paradigm for future applications that probe injury mechanisms and therapeutic interventions for facial paralysis. Our findings provide further evidence contributing to the need to transect both distal branches to ensure facial paralysis in rodents. Despite extensive research on the anatomical contributions of the buccal and marginal mandibular branches to the whisker pad, this is the first study to evaluate these effects at the NMJ level, offering a novel perspective on functional recovery^23,46^. The NMJ is an integral part of determining functional motor recovery following injury^47^. The lack of a standardized model for facial nerve injury and recovery underscores the importance of our findings. By validating functional recovery through histological analysis of target motor endplates, this study provides a more robust framework for assessing therapeutic interventions, thereby contributing to more reproducible and accurate models for facial nerve research.

The lack of innervation of the marginal mandibular in the BB+MM model resulted in significant loss of innervation and loss of NMJ integrity in the whisker pad. These findings were further supported by the assessment of functional motor recovery using the FPS similar to the facial grading systems used for human evaluations^16,43^. This scoring system focused on evaluating the midface of the rat, particularly the nose and whiskers, which are innervated by the buccal and marginal mandibular facial nerves^23,48^. The FPS provides a platform to grossly evaluate recovery over time as the contralateral side remains intact for comparison. Four weeks following injury, the overall FPS for the BB only group was indistinguishable from the sham condition, whereas the BB+MM group remained impaired at 8 weeks following injury.

Following peripheral nerve injury, regenerating axons grow at a rate of 1-2 millimeter per day^49,50^. While the motor axons can regrow, the length and location of the injury impacts their ability to reestablish functional connections with the NMJ, leading to deficits in regaining motor function^51^. In this present model, the distal facial nerves reside 23.76 +/- 1.32 mm from the whisker pad musculature^52^. There is a critical time window of <30 days for the regenerating axons to reach the target muscles and successfully reinnervate the NMJs^51^. Outside of the 30 days window, even if axonal fibers reach the target, they have reduced CMAPs and no motor recovery. Longer denervation times demonstrate no functional recovery and CMAP activity. The lack of compensatory innervation from the marginal mandibular branch in the BB+MM group contributed to the lack of functional recovery. This is particularly poignant given that the axon growth rate and injury site would suggest regrowing axons might reach the muscle as early as 2 weeks post-injury, which is inside the 30-day window for reinnervation. The FPS subscores particularly of the nose asymmetry in the BB+MM condition highlight the importance of the marginal mandibular innervation into the whisker pad, as the lack of whisking could contribute to the lack of function^23,43,46^. This emphasizes the critical role of the marginal mandibular branch in functional recovery and suggests that transecting both nerves is necessary for observing true facial paralysis and assessing functional recovery comprehensively in rodent models of facial paralysis.

We demonstrated that NMJ integrity and morphology aligned with CMAP outputs, with each output confirming that the BB+MM group have prolonged functional deficits beyond 8 weeks. In the presynaptic terminal, shortened axon branches are associated with dystrophic muscles and are a precursor to NMJ collapse^53^. Reduced structural post synaptic changes in the NMJ such as end-plate area, compactness, and AChR cluster dispersion are associated with dystrophic muscles and nerve injury^53–57^. Large AChR area results in larger and stronger CMAPs, which indicate healthy NMJ function ^45,58,59^. Together the pre- and post-synaptic morphological changes observed in the BB+MM transection suggest not only fewer NMJs overall, but an inferior NMJ quality of the remaining structures that would continue to disorganize and disassemble over time^60^. These electrophysiological findings correlate with the observed lack of innervated NMJs and FPS in the double cut model, providing insights into the impact of nerve injury on facial function and potential strategies for assessing and addressing nerve damage in similar contexts.

Our study interrogates the impact on the NMJ following injury and provides several novel findings but has several limitations. There were no considerations for the potential contribution of the autonomic input to the whisker pad, which could contribute to non-facial mediated movement. Additionally, our study focuses on a small animal model, whereas a large animal model would present the opportunity to replicate large gap injuries that mimic those observed in humans. Our model is distal from the main branch, which means that the regenerating nerve has less distance to grow when compared to an injury to the main branch. However, our study is reproducible, provides valuable insight into the NMJ morphology, and introduces a model that will have utility in large gap therapy testing. In the current report, we quantitatively demonstrate the overlap of the buccal and marginal mandibular nerves motor functional recovery by NMJ morphology and innervation. This study expands on the understanding of the interface between the regenerating nerve and the target muscle. The anatomical similarities between the rat facial nerve and humans highlight significant parallels in their structure and function. Early intervention <6 months for facial paralysis is critical for determining patient prognosis^8^. Surgical interventions are the clinical standard of treatment and focus on direct repair of nerve fibers when appropriate^61^. These resemblances provide a valuable model for studying facial nerve regeneration and related medical interventions across species. We have quantitively demonstrated the necessity of transecting both buccal and marginal mandibular nerves to observe complete facial paralysis at the functional and histological level.

## 5. Conclusion

Facial paralysis is a multifaceted condition with broad differential diagnoses and severe impacts on patients’ quality of life. Current management strategies for facial paralysis, while evolving, face significant challenges, particularly in achieving satisfactory functional recovery and addressing complications associated with surgical interventions. The limitations of existing treatments underscore the need for continued development and refinement of therapeutic options to enhance patient outcomes. Rodent models have proven useful in studying facial paralysis and testing potential treatments, but translating these findings to clinical settings has been challenging. This study contributes to the field by demonstrating the impact on the NMJs in the whisker pad of rats following transection both or individual distal branches of the rat facial nerve. These observations provide further understanding about the interface of neural inputs and post synaptic units in the rat whisker pad. By integrating electrophysiological, behavioral, and histological assessments, we offer a comprehensive evaluation of facial nerve injuries that addresses gaps in previous research. Consistent with other investigations, our findings highlight the importance of considering both branches in model development, particularly for studies aiming to understand the compensatory mechanisms and functional recovery of the facial nerve. This approach not only provides a more detailed understanding of nerve injury and recovery but also supports the establishment of standardized models for future research.

## Acknowledgements

This research project was supported through the Office of the Executive Dean for Research at the University of Miami by the Team Neuroscience Award awarded to C.M.D and S.K.G.

## Author Contributions

C.M.D., S.K.G., and B.G.M, conceived and designed the study. B.G.M., J.LG., C.M., G.T., R.R.T., G.M.L., V.L.O., I.G.C., R.M.R., T.R. were responsible for data collection. All authors contributed to the data analysis and interpretation as well as the preparation of the manuscript. All authors contributed to the literature review. All authors have read and agreed to the published version of the manuscript.

## Data Availability Statement

Available data will be provided upon request to the corresponding author.

## Ethics Statement

We confirm that we have read the Journal’s position on issues involved in ethical publication and affirm that this report is consistent with those guidelines.

## Conflicts of Interest

The authors have no conflicts of interest to declare.

## Author Contributions

C.M.D., S.K.G., and B.G.M, conceived and designed the study. B.G.M., J.LG., C.M.,

G.T., R.R.T., G.M.L., V.L.O., I.G.C., R.M.R., T.R. were responsible for data collection. All authors contributed to the data analysis and interpretation as well as the preparation of the manuscript. All authors contributed to the literature review. All authors have read and agreed to the published version of the manuscript.

## Funding

This research was funded by the Team Neuroscience Award awarded by the Office of the Executive Dean for Research.

## Data Availability Statement

Available data will be provided upon request to the corresponding author.

## Conflicts of Interest

The authors have no conflicts of interest to declare.

## Abbreviations

AChR: acetylcholine receptor
BB: buccal transection
BB+MM: buccal + marginal mandibular transection
CMAP: compound muscle action potentials
FPS: Facial palsy score
MEA: microwire electrode array
MM: marginal mandibular
NF200: neurofilament-200
NMJ: Neuromuscular junctions
SV2: synaptic vesicle 2
S100β: S100 calcium-binding protein B
TDT: Tucker-Davis Technologies

## Notes

### Competing Interest Statement

The authors have declared no competing interest.

## References

1. Hohman MH, Hadlock TA. Etiology, diagnosis, and management of facial palsy: 2000 patients at a facial nerve center. The Laryngoscope. 2014;124(7):E283–E293. doi:10.1002/lary.24542

2. Marson AG, Salinas R. Bell’s palsy. West J Med. 2000;173(4):266–268.

3. Mistry RK, Hohman MH, Al-Sayed AA. Facial Nerve Trauma. In: StatPearls. StatPearls Publishing; 2023.

4. Hohman, M.H., Bhama, P.K. and Hadlock, T.A. Epidemiology of iatrogenic facial nerve injury: A decade of experience. The Laryngoscope. 2014;124: 260–265. 10.1002/lary.24117

5. Krane NA, Genther D, Weierich K, et al. Degree of Self-Reported Facial Impairment Correlates with Social Impairment in Individuals with Facial Paralysis and Synkinesis. Facial Plast Surg Aesthetic Med. 2020;22(5):362–369. doi:10.1089/fpsam.2020.0082

6. Nellis JC, Ishii M, Byrne PJ, Boahene KDO, Dey JK, Ishii LE. Association Among Facial Paralysis, Depression, and Quality of Life in Facial Plastic Surgery Patients. JAMA Facial Plast Surg. 2017;19(3):190. doi:10.1001/jamafacial.2016.1462

7. Walker D t., Hallam M j., Ni Mhurchadha S, McCabe P, Nduka C. The psychosocial impact of facial palsy: Our experience in one hundred and twenty six patients. Clin Otolaryngol. 2012;37(6):474–477. doi:10.1111/coa.12026

8. Gordin E, Lee TS, Ducic Y, Arnaoutakis D. Facial Nerve Trauma: Evaluation and Considerations in Management. Craniomaxillofacial Trauma Reconstr. 2015;8(1):1–13. doi:10.1055/s-0034-1372522

9. Sahovaler A, Yeh D, Yoo J. Primary facial reanimation in head and neck cancer. Oral Oncol. 2017;74:171–180. doi:10.1016/j.oraloncology.2017.08.013

10. Razfar A, Lee MK, Massry GG, Azizzadeh B. Facial Paralysis Reconstruction. Otolaryngol Clin North Am. 2016;49(2):459–473. doi:10.1016/j.otc.2015.12.002

11. Ozmen OA, Falcioni M, Lauda L, Sanna M. Outcomes of facial nerve grafting in 155 cases: predictive value of history and preoperative function. Otol Neurotol Off Publ Am Otol Soc Am Neurotol Soc Eur Acad Otol Neurotol. 2011;32(8):1341–1346. doi:10.1097/MAO.0b013e31822e952d

12. Quaranta GC Fabio Piazza, Nicola Quaranta, Ignazio Salonna ,Antonio. Facial Nerve Paralysis in Temporal Bone Fractures: Outcomes after Late Decompression Surgery. Acta Otolaryngol (Stockh). 2001;121(5):652-655. doi:10.1080/00016480119531

13. Mersa B, Tiangco DA, Terzis JK. Efficacy of the. J Reconstr Microsurg. 2000;16:27-36. doi:10.1055/s-2000-7538

14. Bae YC, Zuker RM, Manktelow RT, Wade S. A Comparison of Commissure Excursion following Gracilis Muscle Transplantation for Facial Paralysis Using a Cross-Face Nerve Graft versus the Motor Nerve to the Masseter Nerve. Plast Reconstr Surg. 2006;117(7):2407. doi:10.1097/01.prs.0000218798.95027.21

15. Malik TH, Kelly G, Ahmed A, Saeed SR, Ramsden RT. A Comparison of Surgical Techniques Used in Dynamic Reanimation of the Paralyzed Face. Otol Neurotol. 2005;26(2):284.

16. House, J.W. and Brackmann, D.E. Facial Nerve Grading System. Otolaryngology–Head and Neck Surgery, 1985; 93(2): 146–147. 10.1177/

17. Brown S, Isaacson B, Kutz W, Barnett S, Rozen SM. Facial Nerve Trauma: Clinical Evaluation and Management Strategies. Plast Reconstr Surg. 2019;143(5):1498. doi:10.1097/PRS.0000000000005572

18. Langhals NB, Urbanchek MG, Ray A, Brenner MJ. Update in Facial Nerve Paralysis: Tissue engineering and new technologies. Curr Opin Otolaryngol Head Neck Surg. 2014;22(4):291–299. doi:10.1097/MOO.0000000000000062

19. Höke A. Mechanisms of Disease: what factors limit the success of peripheral nerve regeneration in humans? Nat Clin Pract Neurol. 2006;2(8):448–454. doi:10.1038/ncpneuro0262

20. Ali SA, Stebbins AW, Hanks JE, et al. Facial Nerve Surgery in the Rat Model to Study Axonal Inhibition and Regeneration. J Vis Exp JoVE. 2020;(159). doi:10.3791/59224

21. Hadlock TA, Kowaleski J, Lo D, Mackinnon SE, Heaton JT. Rodent Facial Nerve Recovery After Selected Lesions and Repair Techniques. Plast Reconstr Surg. 2010;125(1):99–109. doi:10.1097/PRS.0b013e3181c2a5ea

22. Chacon MA, Echternacht SR, Leckenby JI. Outcome measures of facial nerve regeneration: A review of murine model systems. Ann Anat - Anat Anz. 2020;227:151410. doi:10.1016/j.aanat.2019.07.011

23. Henstrom D, Hadlock T, Lindsay R, et al. Terminal Segment Surgical Anatomy of the Rat Facial Nerve: Implications for Facial Reanimation Study. Muscle Nerve. 2012;45(5):692–697. doi:10.1002/mus.23232

24. Niimi Y, Matsumine H, Takeuchi Y, et al. A collagen-coated PGA conduit for interpositional-jump graft with end-to-side neurorrhaphy for treating facial nerve paralysis in rat. Microsurgery. 2019;39(1):70–80. doi:10.1002/micr.30291

25. Batioglu-Karaaltin A, Karaaltin MV, Oztel ON, et al. Human olfactory stem cells for injured facial nerve reconstruction in a rat model. Head Neck. 2016;38 Suppl 1:E2011–2020. doi:10.1002/hed.24371

26. Li M, Zhu Q, Liu J. Olfactory ensheathing cells in facial nerve regeneration. Braz J Otorhinolaryngol. 2020;86(5):525–533. doi:10.1016/j.bjorl.2018.07

27. Jang CH, Lee H, Kim M, Kim G. Effect of polycaprolactone/collagen/hUCS microfiber nerve conduit on facial nerve regeneration. Int J Biol Macromol. 2016;93(Pt B):1575–1582. doi:10.1016/j.ijbiomac.2016.04.031

28. Tan J, Xu Y, Han F, Ye X. Genetical modification on adipose-derived stem cells facilitates facial nerve regeneration. Aging. 2019;11(3):908–920. doi:10.18632/aging.101790

29. Fujimaki H, Matsumine H, Osaki H, et al. Dedifferentiated fat cells in polyglycolic acid-collagen nerve conduits promote rat facial nerve regeneration. Regen Ther. 2019;11:240–248. doi:10.1016/j.reth.2019.08.004

30. Zhang Q, Nguyen PD, Shi S, Burrell JC, Cullen DK, Le AD. 3D bio-printed scaffold-free nerve constructs with human gingiva-derived mesenchymal stem cells promote rat facial nerve regeneration. Sci Rep. 2018;8(1):6634. doi:10.1038/s41598-018-24888-w

31. Bengur FB, Komatsu C, Fedor CN, et al. Biodegradable Nerve Guide with Glial Cell Line–Derived Neurotrophic Factor Improves Recovery After Facial Nerve Injury in Rats. Facial Plast Surg Aesthetic Med. Published online March 6, 2023. doi:10.1089/fpsam.2022.0346

32. Tereshenko V, Dotzauer DC, Maierhofer U, et al. Selective Denervation of the Facial Dermato-Muscular Complex in the Rat: Experimental Model and Anatomical Basis. Front Neuroanat. 2021;15. doi:10.3389/fnana.2021.650761

33. Henstrom D, Hadlock T, Lindsay R, et al. The convergence of facial nerve branches providing whisker pad motor supply in rats: Implications for facial reanimation study. Muscle Nerve. 2012;45(5):692–697. doi:10.1002/mus.23232

34. Bueno CR de S, Tonin MCC, Buchaim DV, et al. Morphofunctional Improvement of the Facial Nerve and Muscles with Repair Using Heterologous Fibrin Biopolymer and Photobiomodulation. Pharmaceuticals. 2023;16(5):653. doi:10.3390/ph16050653

35. Greene JJ, McClendon MT, Stephanopoulos N, Álvarez Z, Stupp SI, Richter CP. Electrophysiological assessment of a peptide amphiphile nanofiber nerve graft for facial nerve repair. J Tissue Eng Regen Med. 2018;12(6):1389–1401. doi:10.1002/term.2669

36. Ma F, Zhu T, Xu F, et al. Neural stem/progenitor cells on collagen with anchored basic fibroblast growth factor as potential natural nerve conduits for facial nerve regeneration. Acta Biomater. 2017;50:188–197. doi:10.1016/j.actbio.2016.11.064

37. Matsumine H, Sasaki R, Tabata Y, et al. Facial nerve regeneration using basic fibroblast growth factor-impregnated gelatin microspheres in a rat model. J Tissue Eng Regen Med. 2016;10(10):E559–E567. doi:10.1002/term.1884

38. Matsumine H, Sasaki R, Yamato M, Okano T, Sakurai H. A polylactic acid non-woven nerve conduit for facial nerve regeneration in rats. J Tissue Eng Regen Med. 2014;8(6):454–462. doi:10.1002/term.1540

39. Zhou L, Du HD, Tian HB, Li C, Tian J, Jiang JJ. Experimental study on repair of the facial nerve with Schwann cells transfected with GDNF genes and PLGA conduits. Acta Otolaryngol (Stockh). 2008;128(11):1266–1272. doi:10.1080/00016480801935517

40. Sasaki R, Aoki S, Yamato M, et al. PLGA artificial nerve conduits with dental pulp cells promote facial nerve regeneration. J Tissue Eng Regen Med. 2011;5(10):823–830. doi:10.1002/term.387

41. Karlidag T, Yildiz M, Yalcin S, Colakoglu N, Kaygusuz I, Sapmaz E. Evaluation of the effect of methylprednisolone and N-acetylcystein on anastomotic degeneration and regeneraton of the facial nerve. Auris Nasus Larynx. 2012;39(2):145–150. doi:10.1016/j.anl.2011.03.004

42. Yildirim MA, Karlidag T, Akpolat N, et al. The Effect of Methylprednisolone on Facial Nerve Paralysis With Different Etiologies. J Craniofac Surg. 2015;26(3):810–815. doi:10.1097/SCS.0000000000001502

43. Sasaki R, Matsumine H, Watanabe Y, et al. Electrophysiologic and Functional Evaluations of Regenerated Facial Nerve Defects with a Tube Containing Dental Pulp Cells in Rats. Plast Reconstr Surg. 2014;134(5):970. doi:10.1097/PRS.0000000000000602

44. Takeuchi Y, Osaki H, Matsumine H, Niimi Y, Sasaki R, Miyata M. A method package for electrophysiological evaluation of reconstructed or regenerated facial nerves in rodents. MethodsX. 2018;5:283–298. doi:10.1016/j.mex.2018.03.007

45. Jones RA, Reich CD, Dissanayake KN, et al. NMJ-morph reveals principal components of synaptic morphology influencing structure–function relationships at the neuromuscular junction. Open Biol. 2016;6(12):160240. doi:10.1098/rsob.160240

46. Mattox DE, Felix H. Surgical anatomy of the rat facial nerve. Am J Otol. 1987;8(1):43–47.

47. Bloch-Gallego E. Mechanisms controlling neuromuscular junction stability. Cell Mol Life Sci CMLS. 2015;72(6):1029–1043. doi:10.1007/s00018-014-1768-z

48. Semba K, Egger MD. The facial “motor” nerve of the rat: control of vibrissal movement and examination of motor and sensory components. J Comp Neurol. 1986;247(2):144–158. doi:10.1002/cne.902470203

49. Sulaiman W, Gordon T. Neurobiology of Peripheral Nerve Injury, Regeneration, and Functional Recovery: From Bench Top Research to Bedside Application. Ochsner J. 2013;13(1):100–108.

50. Griffin JW, Pan B, Polley MA, Hoffman PN, Farah MH. Measuring nerve regeneration in the mouse. Exp Neurol. 2010;223(1):60–71. doi:10.1016/j.expneurol.2009.12.033

51. Sakuma M, Gorski G, Sheu SH, et al. Lack of motor recovery after prolonged denervation of the neuromuscular junction is not due to regenerative failure. Eur J Neurosci. 2016;43(3):451–462. doi:10.1111/ejn.13059

52. Pinto MMR, Santos DRD, Bentes LGB, et al. Anatomical description of the extratemporal facial nerve under high-definition system: a microsurgical study in rats. Acta Cir Bras. 2022;37(8): e370803. doi:10.1590/acb370803

53. Pratt SJ, Shah SB, Ward CW, Inacio MP, Stains JP, Lovering RM. Effects of in vivo injury on the neuromuscular junction in healthy and dystrophic muscles. J Physiol. 2013;591(Pt 2):559–570. doi:10.1113/jphysiol.2012.241679

54. Degradation rate of acetylcholine receptors inserted into denervated vertebrate neuromuscular junctions. J Cell Biol. 1989;108(2):647-651.

55. Labovitz SS, Robbins N, Fahim MA. Endplate topography of denervated and disused rat neuromuscular junctions: Comparison by scanning and light microscopy. Neuroscience. 1984;11(4):963–971. doi:10.1016/0306-4522(84)90207-0

56. Sieck DC, Zhan WZ, Fang YH, Ermilov LG, Sieck GC, Mantilla CB. Structure– activity relationships in rodent diaphragm muscle fibers vs. neuromuscular junctions. Respir Physiol Neurobiol. 2012;180(1):88–96. doi:10.1016/j.resp.2011.10.015

57. L L, H Y, H K, et al. Remnant neuromuscular junctions in denervated muscles contribute to functional recovery in delayed peripheral nerve repair. Neural Regen Res. 2020;15(4):731–738. doi:10.4103/1673-5374.266925

58. Hutchinson DO, Walls TJ, Nakano S, Camp S, et al. Congential Endplate acetylocholinesterase deficiency, Brain. 1993; 116(3): 633–653. 10.1093/brain/116.3.633

59. Slater CR, Lyons PR, Walls TJ, Fawcett PR, Young C. Structure and function of neuromuscular junctions in the vastus lateralis of man. A motor point biopsy study of two groups of patients. Brain. 1992; 115(2): 451–479.

60. Burns AS, Jawaid S, Zhong H, et al. Paralysis elicited by spinal cord injury evokes selective disassembly of neuromuscular synapses with and without terminal sprouting in ankle flexors of the adult rat. J Comp Neurol. 2007;500(1):116–133. doi:10.1002/cne.21143

61. Huang H, Lin Q, Rui X, et al. Research status of facial nerve repair. Regen Ther. 2023;24:507–514. doi:10.1016/j.reth.2023.09.012

